## Supplemental Information for "Spatial and functional arrangement of Ebola virus polymerase inside phase-separated viral factories"

**Supplementary Information**

**Supplementary Figure 1: Activity and intracellular localization of the mNG-tagged EBOV VP35. (Related to Figure 1)**

**a** The protein construct of HA-mNG-VP35 and activity of mNG-tagged VP35 (HA-mNG-VP35) relative to the wild-type VP35 (VP35-WT) control measured in supporting the expression of *Renilla* luciferase (Rluc) in the bicistronic EBOV Replication competent minigenome (MG) system. Background expression of MG assessed by excluding L in the MG system (no L). Results from 3 biological replicates with technical triplicates are shown as individual data points with mean ± SD (error bars). **, $p$=0.0005, (N=6, two-tailed, unpaired t tests with Welch's correction). Expression of HA-mNG-VP35 compared to VP35-WT by western blot using a mouse monoclonal ani-VP35 antibody, respectively. Asterisk represents the $\beta$-actin as the loading control.

**b** Confocal immunofluorescence microscopy of fixed HEK 293T cells co-expressing EBOV NP, VP35 (upper panel) or cells co-expressing EBOV NP, VP35 and L (lower panel) at 1 d post-transfection. EBOV NP was labeled with a human monoclonal anti-NP antibody paired with Alexa-568 anti-human antibody. EBOV L was labeled with a rabbit polyclonal anti-L antibody paired with Alexa-568 anti-rabbit antibody. Both VP35 and HA-mNG-VP35 were labeled with a mouse monoclonal anti-FLAG antibody paired with Alexa-647 anti-mouse antibody. Nuclei are counterstained with Hoechst. Representative results from 2 biological replicates with > 3 fields of view are shown. Scale bar: 10 μm.

**c** Size distribution of intracellular NP-VP35 binary EBOV viral factories (VFs) that were used in FRAP experiments. Results from 2 biological replicates with > 3 fields of view (each contains N >100 cells) are analyzed and shown as mean ± SD (error bars).

**d** Non-specific proteins labeled by the rabbit polyclonal anti-L antibody in western blot analysis. Cell lysates collected from the no L control and from VP35-WT (with wild-type L present) sample in (**a**) were analyzed.

**Supplementary Figure 2: Additional characterization of intracellular reconstituted EBOV viral factories. (Related to Figure 1)**

**a** Live-cell imaging of fusion and fission of mNG-labeled, reconstituted EBOV viral factories (VFs). A representative montage of a region in cells co-expressing EBOV NP, HA-mNG-VP35 and L was shown. Similar events were also observed in cells co-expressing EBOV NP and HA-mNG-VP35. White arrows: VF(s) before fusion and fission. Yellow arrows: VF(s) after fusion and fission. Scale bar: 2 μm. Time stamp: (mm:ss).

**b** Trafficking of a non-fluorescent object through the reconstituted EBOV VFs in live cells. A representative montage of a region in cells co-expressing EBOV NP and HA-mNG-VP35 was shown. Similar events were also observed in cells co-expressing EBOV NP, HA-mNG-VP35 and L. Arrows in white indicate the location of the trafficking object. Scale bar: 2 μm. Time stamp: (mm:ss).

**c** Fluorescence recovery of mNG-HA-VP35 in internal reference-regions within VP35-containing condensates. Normalized intensity corresponding to each time point before and after photobleaching of VP35+NP condensate is shown in red, VP35+NP+L condensate is shown in blue. Each data point represents the mean with standard deviation (error bars) of N=9/9 (VP35+NP/VP35+NP+L condensates). Fluorescence intensity was corrected for imaging-induced photobleaching.

**Supplementary Figure 3: Comparative analysis on the internal exchange dynamics of EBOV NP-VP35 binary condensates upon removal of NP disordered region (Related to Figure 1)**

**a** Domain organization of EBOV VP35 and NP. For VP35, 1-80: intrinsically disordered region, binds NP monomer; 80-145: oligomerization domain (OD); 221-340: interferon inhibitory domain

(IID), binds dsRNA. For NP,1-450: N-terminal domain, mediates NP oligomerization and genome encapsidation (ssRNA binding); 450-641: intrinsically disordered region; 641-739: C-terminal domain. NP interacts with VP35 through two independent sites NP (240-285) with VP35(20-48) and NP (481-500)-VP35-IID.

**b** Confocal microscopy of NPΔ (500-739) +VP35 condensates inside live HEK 293T cells 1 day post-transfection. A representative cell from 2 biological replicates (N=9 cells) is shown. The cell body and nucleus (Nuc.) are marked by a dashed line. White arrow: individual condensate chosen for photobleaching. Image montage is composed of selected frames (including t = 0 s) with an interval of 2.75 s from each time-lapse of photobleached condensate displayed. The diameter of photobleached region is 1 μm. Photobleaching occurred at t = 5 s. Scale bars: 5 μm.

**c** Fluorescence recovery of mNG-HA-VP35 within the photobleached region inside intracellular condensates containing VP35. Normalized intensity corresponding to each time point before and after photobleaching of NP (full-length) +VP35 condensate is shown in red, NPΔ (500-739) +VP35 condensate is shown in pink. Each data point represents the mean with standard deviation (error bars) of N=9. Data points with t > 6 s in red and in pink were used to fit a corresponding two-phase association curve, with the normalized intensity value in expressed as a percentage and the curve plateau marked on the side. Goodness of fit of each curve is indicated by an $R^2$ value. For each fitted curve, the best-fit value for each kinetic parameter is shown with a 95% confidence interval.

**Supplementary Figure 4: A reference Vero-VP30 cell infected with the EBOV-GFP-ΔVP30 virus and contains intracellular viral nucleocapsids. (Related to Figure 4)**

**a** Confocal immunofluorescence microscopy of representative Vero-VP30 cells infected with EBOV-GFP-ΔVP30 at MOI (multiplicity of infection) = 3, 18 h post infection. A total of N=39 cells from 2 biological replicates were analyzed. EBOV NP was labeled with a human monoclonal anti-NP antibody paired with Alexa-568 anti-human antibody. EBOV VP30 was labeled with a rabbit monoclonal anti-VP30 antibody paired with Alexa-647 anti-rabbit antibody. Nucleus counterstained with Hoechst. Scale bars: 5 μm.

**b** A subcellular region containing assembled EBOV nucleocapsids appeared as 1 μm-filaments is magnified in single channel view (NP) with a solid red outline. One representative filament was aligned in parallel to a 1 μm-reference bar in red. Scale bar in white: 2 μm.

For display purposes, fluorescence signals of the GFP reporter in EBOV-GFP-ΔVP30 is pseudo-colored yellow; the brightness of insects in NP channel is enhanced. Orthogonal projection of confocal z-stacks shown in (**a**) and (**b**).

**Supplementary Figure 5: Electron microscopy of HEK 293T cells expressing sAPEX2-tagged EBOV polymerase and Pol1-MG, control EBOV polymerase and Pol1-MG and untransfected cells. (Related to Figure 5)** HEK 293T cells were transfected with the Pol1-EBOV-MG system containing sAPEX2-tagged EBOV polymerase (L-VP35) complex or control EBOV polymerase (L-WT+ VP35-V5) and were chemically fixed with 2.5% glutaraldehyde and stained with DAB-$OsO_4$ at 2 d post-transfection.

**a** Electron micrographs of another cytoplasmic region of the DAB+ cell indicated in Figure 5c at different magnifications. Pink arrows: sAPEX2 mediated deposition of electron-dense Osmium. Scale bars: 500 nm/200 nm in micrographs with a medium/high magnification.

**b** Transmission light microscopy (upper) and electron micrograph (lower) of HEK 293T cells transfected with the control EBOV polymerase (L-WT + VP35-V5) and Pol1-EBOV MG systems at 2 days post-transfection. Black arrows indicate electron dense viral factories.

**c** Transmission light microscopy (upper) and electron micrograph (lower) of untransfected HEK 293T cells that were similarly prepared as the DAB+ cells. Scale bars: 20 μm/50 μm in light microscopy images/electron micrographs.

**Supplementary Movie 1: An animated movie sequentially showing tomographic slices and 3D segmentation for the electron tomogram of a subcellular volume, which contains cellular organelles, EBOV viral factories, and sAPEX2-tagged EBOV polymerase. (Related to Figure 6 a, 6b).**

### Supplementary figure 1

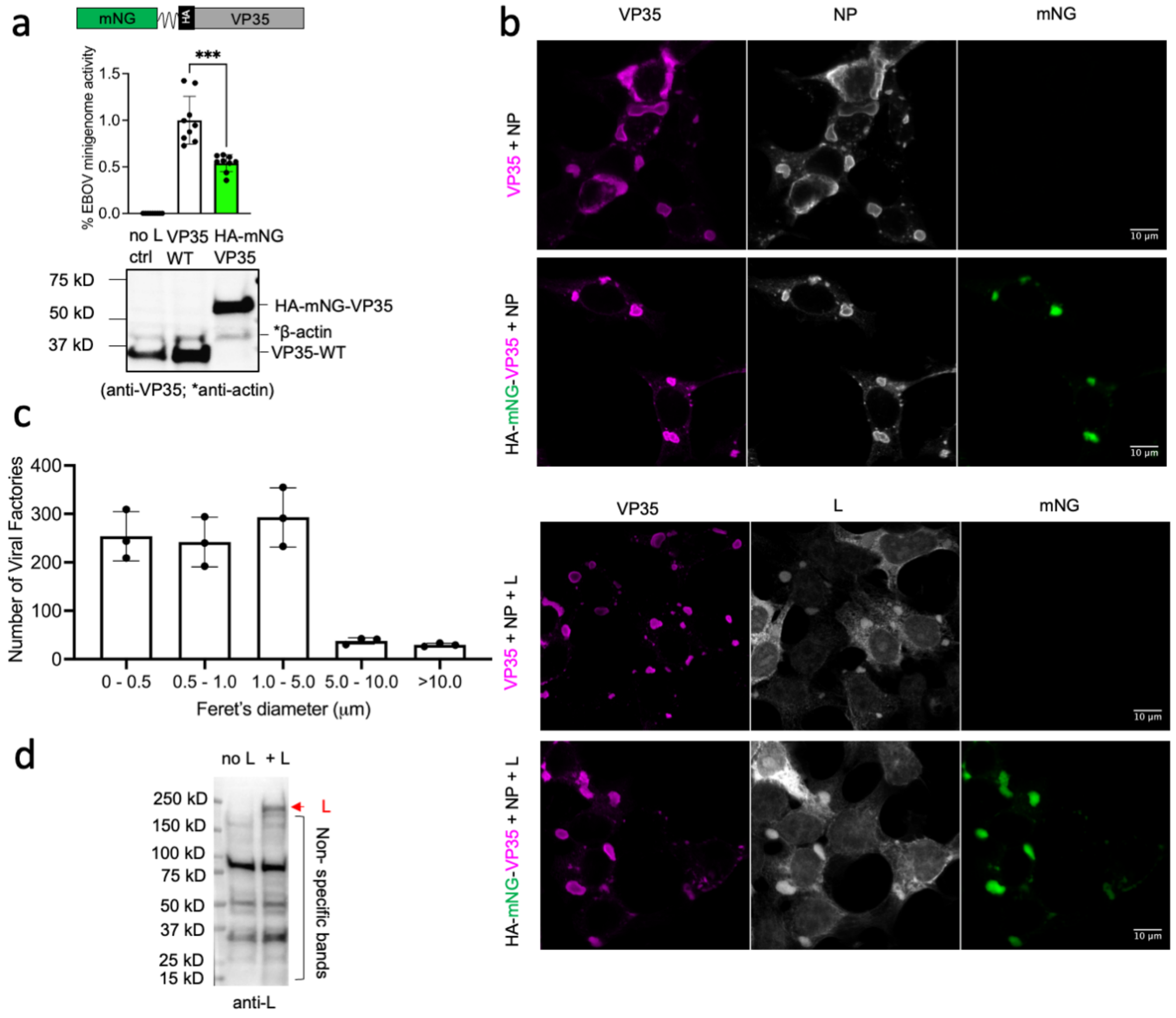

### Supplementary figure 2

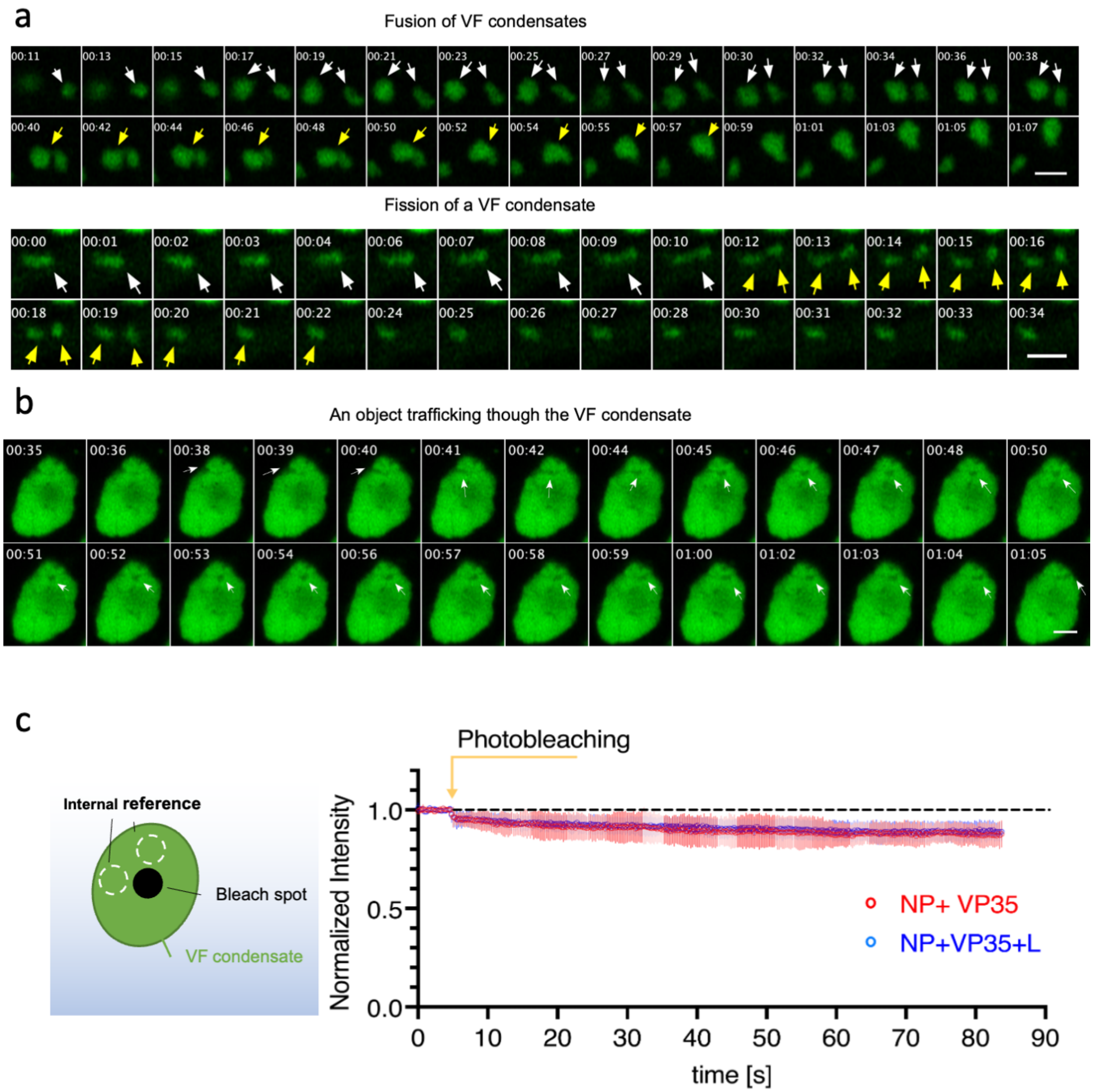

### Supplementary figure 3

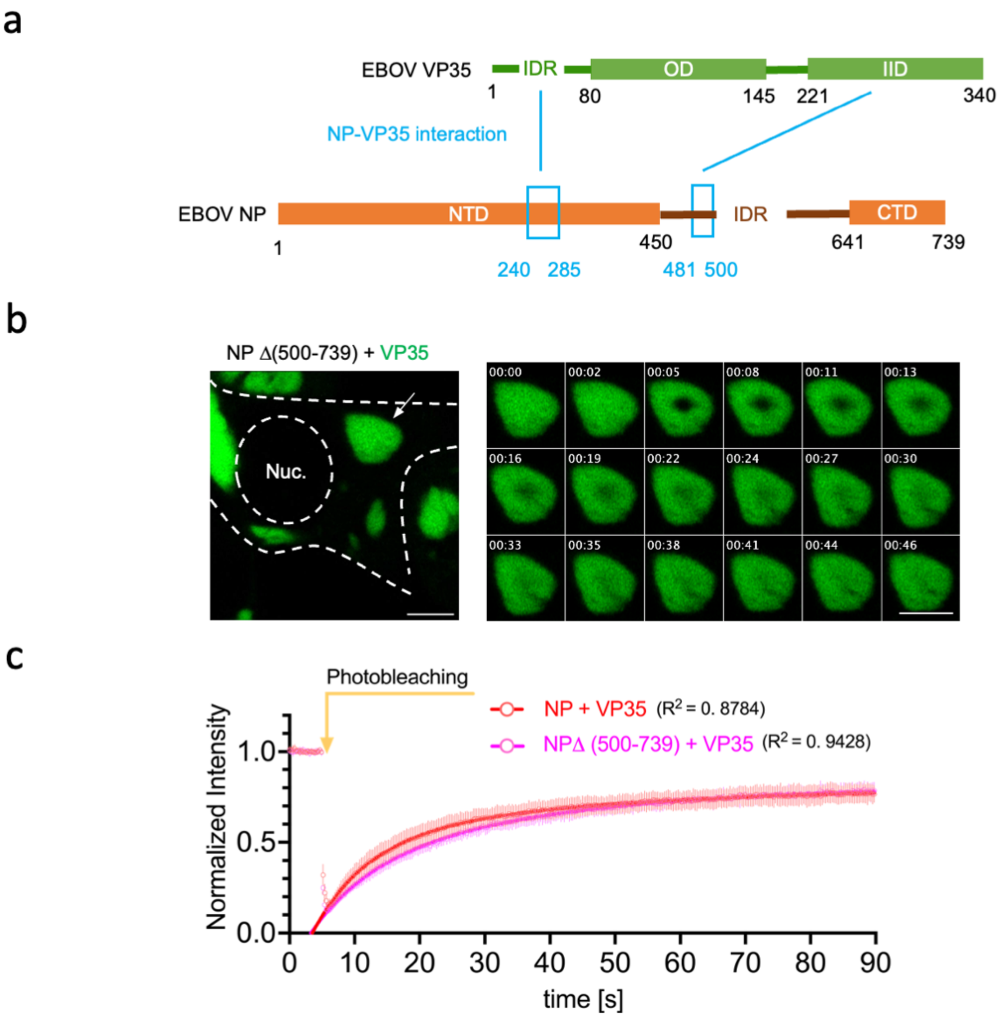


| Fitted parameters | NP + VP35 | NPΔ (500-739) +VP35 |
|---|---|---|
| $t_{1/2}$fast (s) | 5.799 (4.341, 7.258) | 6.313 (4.231, 8.970) |
| $t_{1/2}$slow (s) | 26.22 (17.80, 76.37) | 22.95 (18.52, 43.25) |
| %fast | 69.52 (58.73, 76.37) | 47.60 (36.70, 65.51) |

### Supplementary figure 4

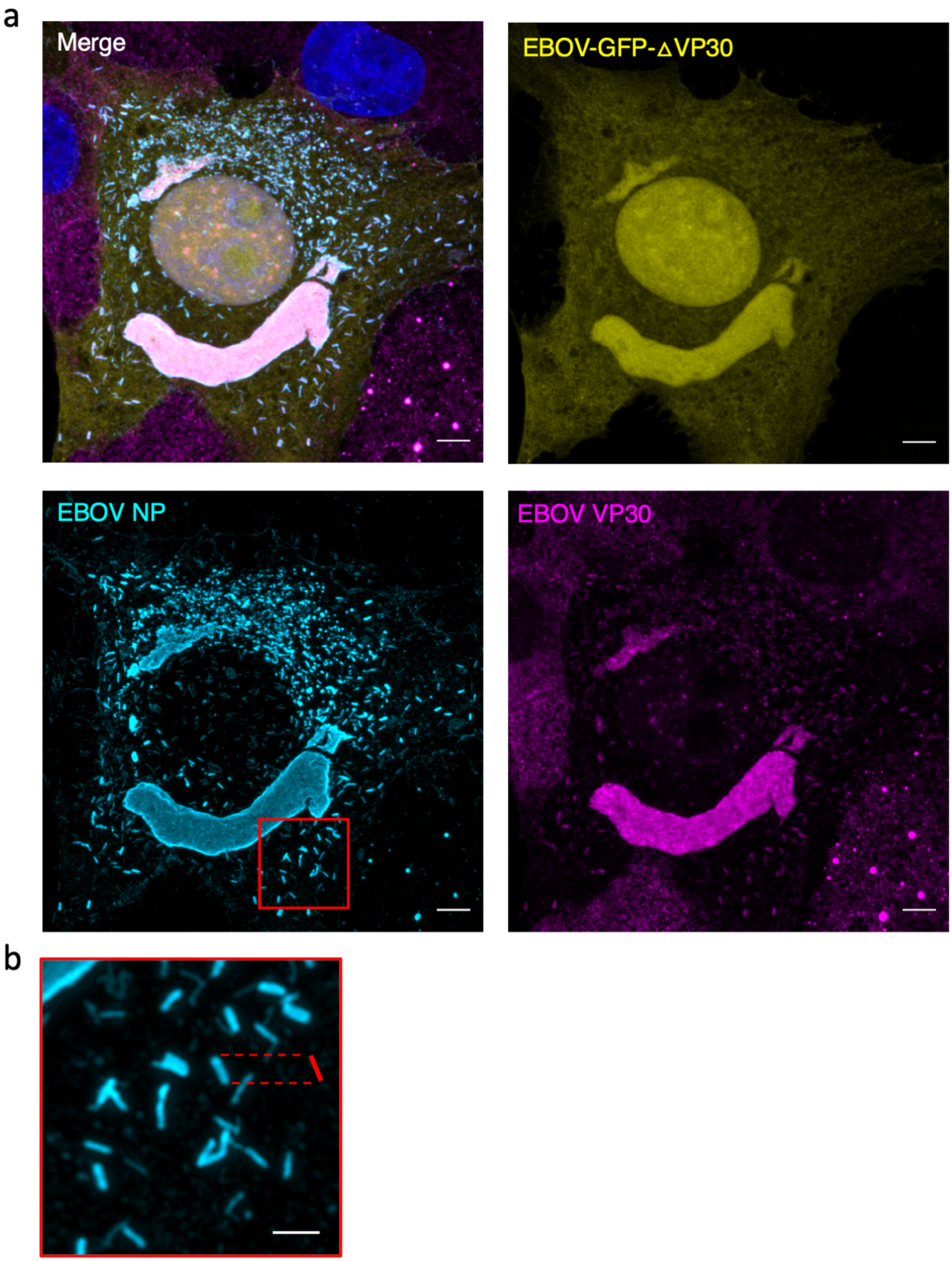

### Supplementary figure 5

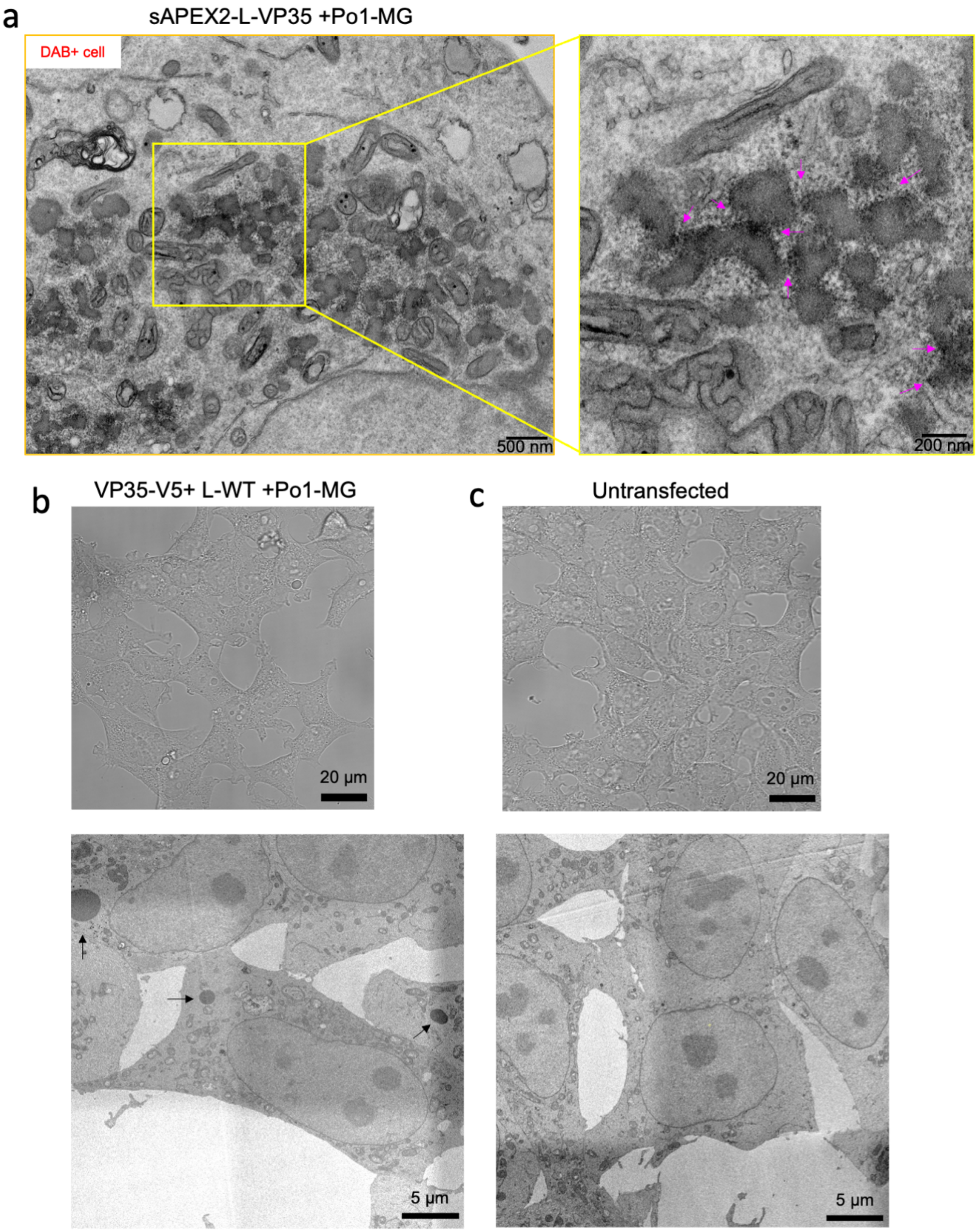
